# Quantitative multidimensional phenotypes improve genetic analysis of laterality traits

**DOI:** 10.1101/2021.04.28.441754

**Authors:** Judith Schmitz, Mo Zheng, Kelvin F. H. Lui, Catherine McBride, Connie S.-H. Ho, Silvia Paracchini

## Abstract

Handedness is the most commonly investigated lateralised phenotype and is usually measured as a binary left/right category. Its links with psychiatric and neurodevelopmental disorders prompted studies aimed at understanding the underlying genetics, while other measures and side preferences have been less studied. We investigated the heritability of hand, as well as foot, and eye preference by assessing parental effects (*n* ≤ 5 028 family trios) and SNP-based heritability (SNP-h^2^, *n* ≤ 5 931 children) in the Avon Longitudinal Study of Parents and Children (ALSPAC). An independent twin cohort from Hong Kong (*n* = 358) was used to replicate results from structural equation modelling (SEM). Parental left-side preference increased the chance of an individual to be left-sided for the same trait, with stronger maternal than paternal effects for footedness. By regressing out the effects of sex, age, and ancestry, we transformed laterality categories into quantitative measures. The SNP-h^2^ for quantitative handedness and footedness was .21 and .23, respectively, which is higher than the SNP-h^2^ reported in larger genetic studies using binary handedness measures. The heritability of the quantitative measure of handedness increased (.45) compared to a binary measure for writing hand (.27) in the Hong Kong twins. Genomic and behavioural SEM identified a shared genetic factor contributing to handedness, footedness, and eyedness, but no independent effects on individual phenotypes. Our analysis demonstrates how quantitative multidimensional laterality phenotypes are better suited to capture the underlying genetics than binary traits.

## Introduction

The cerebral hemispheres differ in function and structure underpinning specialisation for cognition, perception, and motor control ^1^. For instance, language is predominantly processed in the left hemisphere in most individuals ^2^ and the *planum temporale* typically shows a pronounced structural leftward asymmetry ^3^, although there is little evidence for a strong association between the two forms of asymmetry ^4^. Neurodevelopmental disorders such as dyslexia ^5,6^, schizophrenia ^7^, or autism spectrum disorder (ASD) ^8^ have been associated with a higher prevalence of atypical *planum temporale* asymmetry.

The most commonly studied lateralised trait is handedness. Worldwide, around 10% of the general population is left-handed with slight geographical variation ^9^, likely influenced by cultural factors ^10,11^. Meta-analyses have confirmed higher rates of left- or non-right-handedness in ASD ^12^ and schizophrenia ^13^. A genetic influence on handedness has been inferred from family and adoption studies ^14^. For instance, the probability of left-handedness increases with the number of left-handed parents ^15^. Twin studies reported slightly higher rates of concordance in monozygotic (MZ) compared to dizygotic (DZ) twins ^16,17^ and provided heritability estimates of around .25 ^18,19^.

Family studies have suggested differential effects of fathers and mothers to their offspring’s handedness. A stronger maternal than paternal effect was repeatedly found in biologically related parent-offspring trios ^20,21^ and a similar trend was observable in an adoption study ^22^. A maternal effect on non-right-handedness was also found in 592 families, where a paternal effect was only detectable in males ^23^.

A recent large-scale genome-wide association study (GWAS; *n* ∼ 2M) estimated that up to 6% of the variance in left-handedness and up to 15% of the variance in ambidexterity are explained by common genetic markers ^24^. As in most large-scale laterality studies, handedness was assessed as hand preference for writing, leading to three categories: right, left or both. The “both” category identifies individuals who say that they can write equally well with both hands, referred to as ambidextrous. However, a single task cannot identify mixed-handed individuals who prefer different hands for different activities. Instead, self-report questionnaires such as the Edinburgh Handedness Inventory (EHI) ^25^ assess the preferred hand for several manual activities and therefore capture both mixed-handed and ambidextrous individuals. A GWAS on brain imaging parameters (*n* = 32 256) revealed that genetic markers associated with structural brain asymmetries overlapped with markers previously associated with writing hand preference. Moreover, genetic factors involved in brain asymmetry overlap with neurodevelopmental and cognitive traits such as ASD, schizophrenia, educational attainment (EA) ^26^, and intelligence (IQ) ^27^. These data suggest a general mechanism for the establishment of left/right asymmetry which is also important for neurodevelopmental outcomes. Therefore, the analysis of other lateralised preferences will contribute to the understanding of such general mechanisms.

Foot and eye preference have received considerably less attention, even though associations with neurodevelopmental disorders have been reported as well. For example, we found an increased prevalence of non-right-footedness in neurodevelopmental and psychiatric disorders (*n*_cases_ = 2 431, *n*_controls_ = 116 938) ^28^. Smaller studies point to higher rates of left eye preference in schizophrenia (*n*_cases_ = 88, *n*_controls_ = 118 ^29^; *n*_cases_ = 68, *n*_controls_ = 944 ^30^) and ASD (*n*_cases_ = 37; *n*_controls_ = 20) ^31^. Warren et al. ^32^ reported heritability estimates for foot and eye preference to be .12 and .13, respectively. In Japanese twins, Suzuki and Ando ^33^ provided heritability estimates for foot preference ranging from .08 to .24 and having one left-footed parent increased the probability of being left-footed ^34^. These studies support a genetic component for foot and eye preference although there is variability in heritability estimates, probably resulting from small sample sizes.

We performed the largest heritability study to date for multiple side preferences in the Avon Longitudinal Study of Parents and Children (ALSPAC) and a twin cohort from Hong Kong to investigate the heritability of laterality phenotypes, their associations with one another, and their links to neurodevelopmental and cognitive outcomes.

## Materials and Methods

### Cohorts

#### ALSPAC

ALSPAC is a population-based longitudinal cohort. Pregnant women living in Avon, UK, with expected dates of delivery from 1st April 1991 to 31st December 1992 were invited to take part, resulting in 14 062 live births and 13 988 children who were alive at 1 year of age ^35,36^. Informed consent for the use of data collected via questionnaires and clinics was obtained from participants following the recommendations of the ALSPAC Ethics and Law Committee at the time. Ethical approval for the study was obtained from the ALSPAC Ethics and Law Committee and the Local Research Ethics Committees. Please note that the study website contains details of all the data that is available through a fully searchable data dictionary and variable search tool (http://www.bristol.ac.uk/alspac/researchers/our-data/).

#### Hong Kong

Study participants were recruited from the Chinese-English Twin Study of Biliteracy, a longitudinal study of primary school twin children starting in 2014 ^37^. Participating children were recruited from Hong Kong primary schools and had Cantonese as their native language. Language and cognitive ability tests have been conducted for over four waves with a one-year interval between assessments. Laterality data were collected during the second wave of assessment.

### Participants and phenotypes

#### ALSPAC

Laterality phenotypes were assessed for children based on maternal reports and for parents as self-report. Hand preference was assessed using eleven items for parents and six items for children. Foot preference and eye preference were assessed using four and two items, respectively, for parents and children. All items were rated on a 3-point scale (coded as left = 1, either = 2, right = 3, see Table S1). Two summary items (one in a right-mixed-left [R-M-L] classification and one in a right-left classification [R-L]) were derived from recoded mean values across non-missing items for hand, foot, and eye preference (see supplementary methods and Figures S1-S3 for details). Mean ages of mothers, fathers, and children were 32.54 (SD = 4.42), 34.42 (SD = 5.60) and 3.55 (SD = 0.07) at the time of assessment, respectively.

#### Hong Kong

The overall sample comprised *n* = 366 twin children (183 twin pairs) with a mean age of 8.67 years (SD = 1.23). This sample included 81 MZ pairs (37 male pairs and 44 female pairs) and 102 DZ pairs (21 male pairs, 19 female pairs, and 62 opposite-sex pairs). Twin zygosity of same-sex twins was determined by genotyping small tandem repeat (STR) markers on chromosomes 13, 18, 21, X and Y by Quantitative Fluorescence-Polymerase Chain Reaction (QF-PCR).

Hand, foot, and eye preference were assessed using a modification of the EHI ^25^. The questionnaire was translated into Chinese and included six hand preference items, one foot preference item, and one eye preference item. All items were read to participants by a trained research assistant as described in detail previously ^38^. Items were coded to a 3-point scale and a R-M-L summary item was created for hand preference (see supplementary methods and Figure S4 for details).

### Genotype quality control (QC)

#### ALSPAC

Children’s genotypes were generated on the Illumina HumanHap550-quad array at the Wellcome Trust Sanger Institute, Cambridge, UK and the Laboratory Corporation of America, Burlington, NC, US. Standard QC was performed as described elsewhere ^39^. In total, 9 115 children and 500 527 SNPs passed QC filtering.

#### Hong Kong

Genotyping was performed using Illumina Human Infinium OmniZhongHua-8 v1.3 Beadchip at the Prenatal Genetic Diagnosis Centre and the Pre-implantation Genetic Diagnosis laboratory in the Prince of Wales Hospital and The Chinese University of Hong Kong, Hong Kong SAR. Standard quality control measures were carried out. Genetic variants with missing rate > 10%, minor allele frequency (MAF) < 0.01 and with significant deviation from Hardy-Weinberg equilibrium (*p* < 1 ×10^−6^) were excluded. Individuals with genotyping rates < 90% and outlying heterozygosity rates were excluded. In total, 911 178 SNPs passed QC filtering. Among the *n* = 366 twin children, genotype data were available for *n* = 358 (81 MZ pairs and 98 DZ pairs).

### Parental effects

We included parent-child trios with complete phenotypic data on the summary items for hand, foot or eye preference after excluding one of each twin pair (*n* = 113) and children with physical disabilities (*n* = 65) or sensory impairments (*n* = 50), resulting in a sample size (number of trios) of *n*_hand_ = 5 028, *n*_foot_ = 4 960 and *n*_eye_ = 4 762 (see Table S1).

For hand, foot, and eye preference, we first performed two logistic regression analyses using both parents’ sidedness as a predictor (coded as 0 = two right-sided parents, 1 = one mixed-sided parent, 2 = one left-sided parent, 3 = two mixed-sided parents, 4 = one mixed- and one left-sided parent, 5 = two left-sided parents). This analysis was performed for child sidedness (coded as right = 0, left = 1) using both the A) R-M-L classification (excluding mixed-sided children and their parents) and the B) R-L classification.

Next, we differentiated maternal and paternal effects by using maternal sidedness, paternal sidedness (both coded as right = 0, mixed = 1, left = 2), and offspring sex, as well as interaction terms between maternal and paternal sidedness with offspring sex as predictors. We used the wald.test() function to test for a difference between maternal and paternal effects using the R-M-L and the R-L classification.

As non-paternity could affect these analyses, we reran the logistic regression analyses including only confirmed biological parent-offspring trios as confirmed by genotype data. Genotypes were available for *n* = 1 719 fathers. We used the R package Sequoia ^40^, which assigns parents to offspring based on Mendelian errors. Sequoia uses birth year and sex to decrease the number of potential relationships between individuals and to correctly infer parents and offspring. As the exact birth year of children and parents in ALSPAC was unknown to us, children’s birth year was set to 1992 and parents’ birth year was roughly estimated from the age of the assessment of laterality data. We selected 500 SNPs randomly from a subset that had MAF > 0.45, high genotyping rate (missingness < 0.01) and low linkage disequilibrium (LD; r^2^ < 0.01 within a 50 kb window). The 500 SNPs were spread across chromosomes 1 to 22. Sequoia confirmed paternity for *n* = 1 624 fathers. Among this subsample of 1 624 trios, complete phenotypic data were available for 1 161 trios for handedness, 1 150 trios for footedness, and 1 105 trios for eyedness (see Table S1).

To assess the reliability of maternal reports, we performed Spearman rank correlation analysis between hand preference for drawing (left/right) assessed by maternal report at age 3.5 and self-reported hand preference for writing at age 7.5 (M_age_ = 7.50 years; *n* = 3 129).

### Phenotypic analysis

Unrelated children (genetic relationship < 0.05, *n* = 5 956) with genome-wide genetic and phenotypic data were selected for Genome-wide Complex Trait Analysis (GCTA) ^41^. The same sample was used for phenotypic analysis. Sample sizes varied from *n* = 4 630 (foot used to pick up a pebble) to *n* = 5 931 (summary item for hand preference).

Summary items in the R-M-L classification for hand, foot, and eye preference and 12 single items were residualised for sex, age, and the two most significant principal components:

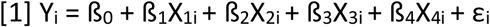

Where Y_i_ is the laterality summary item (coded as right = 0, mixed = 1, left = 2), ß_0_ is the intercept, ß_1_ is the regression weight for offspring sex, X_1i_ is offspring sex, ß_2_ is the regression weight for offspring age in weeks, X_2i_ is offspring age in weeks, ß_3_ is the regression weight for PC1, X_3i_ is PC1, ß_4_ is the regression weight for PC2, X_4i_ is PC2, and ε_i_ reflects random error.

Phenotypes were then inverse rank-transformed to achieve normally distributed phenotypes. Principal components were calculated based on directly genotyped (MAF < 0.05) and LD pruned (r^2^ < 0.01 within a 50 kb window) SNPs (excluding high LD regions) using Plink v2. The rationale for including PCs in the phenotype transformation was based on the Genetic-relationship-matrix structural equation modelling (GRM-SEM) method which has been developed using the ALSPAC cohort ^42^. As there is little population stratification in ALSPAC, the PC effect on the phenotypes is very small. Instead, higher scores indicated being left-sided, being female ^43,44^, and younger age. Phenotypic correlations between rank-transformed items were calculated with Pearson correlation, applying FDR correction for 105 comparisons using the Benjamini Hochberg method ^45^.

### Heritability estimates

SNP-h^2^ was calculated for the transformed R-M-L summary items (3) and single items (12) using restricted maximum-likelihood (REML) analysis in GCTA ^46^, which compares phenotypic similarity and genotypic similarity based on a genetic-relationship matrix (GRM) in unrelated individuals. A GRM was estimated based on directly genotyped SNPs for unrelated children (genetic relationship < 0.05, *n* = 5 956) using GCTA.

As a comparison, SNP-h^2^ was calculated for the untransformed categorical items using sex, age, and the first two principal components as covariates. We estimated SNP-h^2^ separately for left-sidedness (left vs. right, excluding mixed-sided individuals) and mixed-sidedness (mixed vs. right, excluding left-sided individuals).

Next, we estimated heritability from parent-offspring data ^47^. Among the subsample with genomic data and confirmed paternity, we selected those with information on age at the time of laterality assessment, resulting in a sample of 1 000 trios for handedness, 991 trios for footedness, and 957 trios for eyedness. Summary items in the R-M-L classification for hand, foot, and eye preference (coded as right = 0, mixed = 1, left = 2) were transformed following the same procedure described above for the ALSPAC children. We estimated heritability by performing linear regression analyses using mean parental laterality as predictor and child laterality as the outcome:

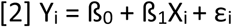

Where Y_i_ is the transformed offspring laterality item, ß_0_ is the intercept, ß_1_ is the regression weight (heritability index), X_i_ is the mean parental laterality, and ε_i_ reflects random error.

### SEM

We applied GRM-SEM ^42^ to quantify shared and unique genetic factors among R-M-L summary items for hand, foot, and eye preference. This method has recently been used to study genetic associations among language and literacy skills in the ALSPAC cohort ^48^. Equivalent to heritability analysis in twin research, GRM-SEM partitions phenotypic variance/covariance into genetic and residual components, but estimates genetic variance/covariance based on genome-wide genetic markers. We used the same GRM described above (based on directly genotyped SNPs for *n* = 5 956 unrelated children using GCTA). A GRM-SEM was fitted using the grmsem library in R (version 1.1.0) using all children with phenotypic data for at least one phenotype. Multivariate trait variances were modelled using a saturated model (Cholesky decomposition). GRM-SEM was also used to estimate bivariate heritability, i.e. the contribution of genetic factors to the phenotypic covariance.

The heritability of laterality phenotypes was additionally estimated using a classical twin design that compares the similarity of MZ to that of DZ twins. Since MZ twins share nearly all their genetic variants, whereas DZ twins share on average 50% of their genetic variants, any excess similarity of MZ twins over DZ twins is the result of genetic influences. This method partitions phenotypic variance into that due to additive genetic (A), shared environmental (C) and non-shared environmental influence (E). The variance attributed to each component can be estimated using the structural equation modelling (SEM) technique and the proportion of variance explained by the genetic influence (A) is termed heritability. Phenotypes were transformed following the same procedure described for ALSPAC above. We fit a multivariate ACE model to the transformed phenotypes (handedness, footedness, and eyedness) and compared ACE with its constrained models, such as the AE model. Analyses were performed using the OpenMx software package 2.18.1 ^49^. The script was adapted from the International Workshop on Statistical Genetic Methods for Human Complex Traits ^50^.

### Polygenic risk score (PRS) analysis

We conducted PRS analyses using summary statistics for handedness assessed as a binary trait, psychiatric and neurodevelopmental conditions (ASD, ADHD, bipolar disorder (BIP), schizophrenia (SCZ)), and cognitive measures (EA and IQ) using PRSice 2.3.3 ^51^. PRS analyses were performed for hand and foot preference (which showed significant SNP-h^2^). The summary statistics for hand preference (left vs. right) were calculated after excluding individuals from 23andMe as well as ALSPAC from the original GWAS ^24^ sample. Summary statistics for ADHD ^52^, ASD ^53^, BIP ^54^, and SCZ ^55^ were accessed from the Psychiatric Genomics Consortium (PGC) website (https://www.med.unc.edu/pgc/data-index/). Summary statistics for IQ ^56^ and EA ^57^ were accessed from the Complex Trait Genetics (CTG) lab website (https://ctg.cncr.nl/software/summary_statistics), and the Social Science Genetic Association Consortium (https://www.thessgac.org/data), respectively.

PRS were derived from LD-clumped SNPs (r^2^ < 0.1 within a 250 kb window) as the weighted sum of risk alleles according to the training GWAS summary statistics. No covariates were included as phenotypes had been corrected for effects of age, sex, and ancestry. Results are presented for the best training GWAS *p*-value threshold (explaining maximum phenotypic variance) as well as GWAS *p*-value thresholds of .001, 0.05, .1, .2, .3, .4, .5, and 1. Results were FDR-corrected for 126 comparisons (7 training GWAS; 2 target phenotypes; 9 *p*-value thresholds) using the Benjamini-Hochberg method ^45^.

### Code availability

Data preparation and visualization were performed using R v.4.0.0. Analysis scripts are available through Github (https://github.com/Judith-Schmitz/heritability_hand_foot_eye).

## Results

### Parental effects

We tested parental effects by assessing the percentages of non-right-sided (R-M-L) and left-sided (R-L) offspring as a function of parental sidedness in the whole sample and in trios with confirmed biological paternity. As expected, the percentage of non-right-sidedness and left-sidedness were highest in individuals with two non-right-sided or two left-sided parents, respectively (Tables S2 [R-M-L] and S3 [R-L]). The percentage of non-right-sidedness and left-sidedness were higher in individuals with a non-right-sided or left-sided mother and a right-sided father than vice versa for all three traits. This effect was visible in both the whole sample (e.g. 31.23% vs. 25.83% for non-right-handedness, see Table S2) and in the subset with confirmed biological paternity (e.g. 33.33% vs. 25.37%, see Table S2).

Second, we ran logistic regression analyses in *n* ≤ 5 028 ALSPAC family trios. In the R-M-L classification (*n*_hand_ = 4 248, *n*_foot_ = 3 242 and *n*_eye_ = 3 050), parental sidedness predicted hand, Х^2^(5) = 39.5, *p* = 1.9 × 10^−7^, foot, Х^2^(5) = 59.9, *p* = 1.3 × 10^−11^, and eye preference, Х^2^(5) = 27.4, *p* = 4.8 × 10^−5^. In the R-L classification (*n*_hand_ = 5 028, *n*_foot_ = 4 960 and *n*_eye_ = 4 762), parental sidedness also predicted hand, Х^2^(2) = 42.6, *p* = 5.5 × 10^−10^, foot, Х^2^(2) = 69.1, *p* = 1.0 × 10^−15^, and eye preference, Х^2^(2) = 14.6, *p* = 6.9 × 10^−4^. ORs show that having one or two left-sided parents increased one’s chances to be left-sided for hand, foot, and eye preference in the R-M-L classification (Figure 1A) and in the R-L classification (Figure 1B). Analysis in the subsample with confirmed paternity (*n* ≤ 1 161 family trios) showed similar, although attenuated, parental effects for hand (R-M-L: Х^2^(4) = 14.9, *p* = .005; R-L: Х^2^(2) = 12.1, *p* = .002) and foot (R-M-L: Х^2^(4) = 22.5, *p* = .0002; R-L: Х^2^(2) = 19.1, *p* = 7.1 × 10^−5^), but not for eye (R-M-L: Х^2^(5) = 5.3, *p* = .380; R-L: Х^2^(2) = 2.7, *p* = .250) preference (Figure S5). The full regression model outputs for the whole sample and for trios with confirmed paternity can be found in Tables S4-S7.

**Figure 1:**
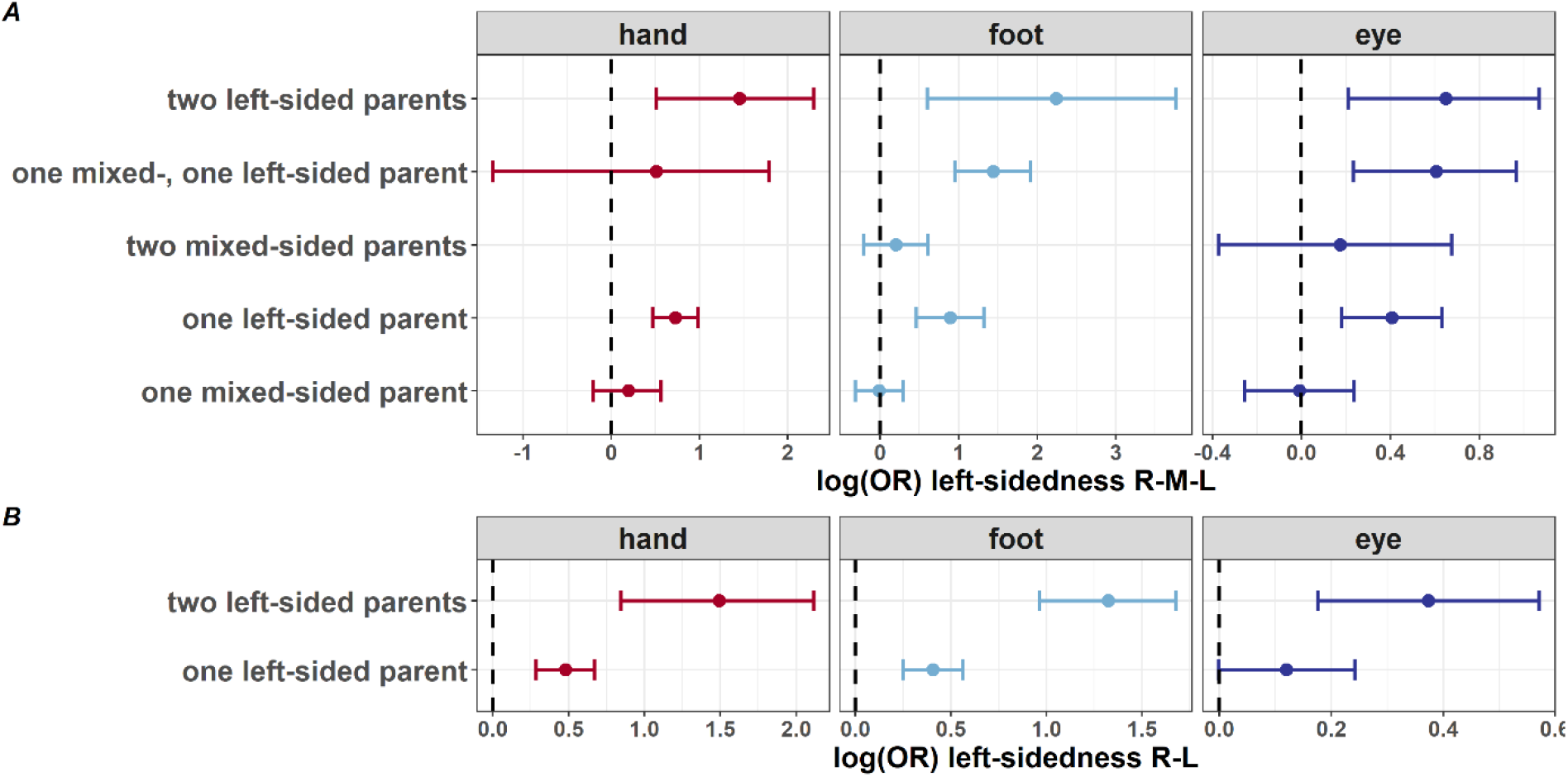
Parental effects on child sidedness. ORs [95% CI], resulting from logistic regression analysis.

Third, we investigated maternal and paternal effects and possible interactions with offspring sex. In the whole sample, Wald tests revealed a significant maternal effect on hand (R-M-L: Х^2^(4) = 38.9, *p* = 7.4 × 10^−8^; R-L: Х^2^(2) = 31.7, *p* = 1.3 × 10^−7^), foot (R-M-L: Х^2^(4) = 52.7, *p* = 9.8 × 10^−11^; R-L: Х^2^(2) = 96.6, *p* < 2.2 × 10^−16^), and eye preference (R-M-L: Х^2^(4) = 38.3, *p* = 9.7 × 10^−8^; R-L: Х^2^(2) = 34.1, *p* = 3.9 × 10^−8^). Paternal sidedness predicted hand (R-M-L: Х^2^(4) = 10.3, *p* = .036; R-L: Х^2^(2) = 12.3, *p* = .002) and foot (R-M-L: Х^2^(4) = 15.1, *p* = .005; R-L: Х^2^(2) = 6.0, *p* = .049), but not eye preference (R-M-L: Х^2^(4) = 4.6, *p* = .330; R-L: Х^2^(2) = 0.6, *p* = .760). Wald tests contrasting maternal and paternal effects revealed a stronger maternal than paternal effect only for foot preference (R-M-L: Х^2^(1) = 4.6, *p* = .033; R-L: Х^2^(1) = 23.9, *p* = 1.0 × 10^−6^). This effect was confirmed in the subsample with confirmed paternity (R-M-L: Х^2^(1) = 8.4, *p* = .004; R-L: Х^2^(1) = 10.0, *p* = .002). Although attenuated in the smaller subsample with confirmed paternity, this finding suggests a genuinely stronger maternal than paternal effect on footedness. In the whole sample, interaction terms between maternal/paternal sidedness and offspring sex revealed that in the R-L classification, maternal left-sidedness had a greater effect on left-footedness in girls compared to boys (ß = 0.49, SE = 0.19, *z* = 2.55, *p* = .011), which was confirmed in the smaller subsample (ß = 0.96, SE = 0.44, *z* = 2.17, *p* = .030). The full regression model outputs for both the whole sample and the subsample with confirmed paternity can be found in Tables S8-S11.

Besides non-paternity, the reliability of the maternal report on laterality phenotypes could have affected our analysis. Correlation analysis showed a strong association between hand preference for drawing collected at 3.5 years of age and the self-reported hand preferred for writing at age 7.5 (*r* = .95, 95% CI = [.93, .97], *p* < 2.2 × 10^−16^). Among the 2 838 children with a right-hand preference at age 3.5, seven reported a left-hand preference for writing at age 7.5. Of the 291 children with left-hand preference at age 3.5, 19 showed a right-hand preference for writing at age 7.5. Overall, 99.2% of individuals showed stable hand preference (see Table S12), demonstrating the reliability of the maternal report.

### Transformed phenotypes

Phenotypic correlation and genomic analyses (SNP-h^2^ estimates, GRM-SEM and PRS analysis) were performed in unrelated children from the ALSPAC cohort (*n* ≤ 5 931). Multivariate behavioural SEM analysis was performed In the Hong Kong twin sample (*n* ≤ 358). The absolute numbers and percentages of children with left, mixed and right side preference for the three summary items in both cohorts are shown in Table 1.

**Table 1:** Children with left, mixed and right side preference for each phenotype in ALSPAC (unrelated children) and the Hong Kong cohort (twin children).

| ALSPAC |  |  |  |  | Hong Kong |  |  |  |
| --- | --- | --- | --- | --- | --- | --- | --- | --- |
|  | *n* | Left | Mixed | Right | *n* | Left | Mixed | Right |
| Hand preference | 5 931 | 471 | 893 | 4567 | 358 | 20 | 37 | 301 |
|  |  | (7.9%) | (15.1%) | (77.0%) |  | (5.6%) | (10.3%) | (84.1%) |
| Foot preference | 5 860 | 344 | 2070 | 3446 | 358 | 31 | 106 | 221 |
|  |  | (5.9%) | (35.3%) | (58.8%) |  | (8.7%) | (29.6%) | (61.7%) |
| Eye preference | 5 650 | 730 | 2012 | 2908 | 357 | 95 | 107 | 155 |
|  |  | (12.9%) | (35.6%) | (51.5%) |  | (26.5%) | (29.9%) | (43.4%) |

By regressing out the effects of sex, age, and ancestry, we transformed laterality categories into quantitative measures using formula [1]. We assessed phenotypic correlations for the transformed items in ALSPAC and the Hong Kong cohort. In ALSPAC, the single item that best captured the summary item was “hand used to draw” for hand preference (*r* = .87, *t*_(5920)_ = 139.01, *p* < 2.2 × 10^−16^), “foot used to stamp” for foot preference (*r* = .78, *t*_(5765)_ = 95.78, *p* < 2.2 × 10^−16^), and “eye used to look through a bottle” for eye preference (*r* = .96, *t*_(5469)_ = 249.61, *p* < 2.2 × 10^−16^). In both cohorts, summary items showed positive correlations with each other (Figure S6, Figure S7). These correlations support a general left/right directionality captured by the different items.

### Heritability estimates

We then tested the heritability of the transformed phenotypes. SNP-h^2^ of transformed laterality items ranged from .00 (*p* = .500) for “eye used to look through a bottle” to .42 (*p* = 8 × 10^−13^) for “hand used to cut” (Figure 2, Table S13). The highest heritability estimate for summary measures was observed for footedness (SNP-h^2^ = .23; *p* = 2 × 10^−5^), followed by handedness (SNP-h^2^ = .21; *p* = 1 × 10^−4^). There was no significant SNP-h^2^ for eyedness (SNP-h^2^ = .00; *p* = .469).

**Figure 2:**
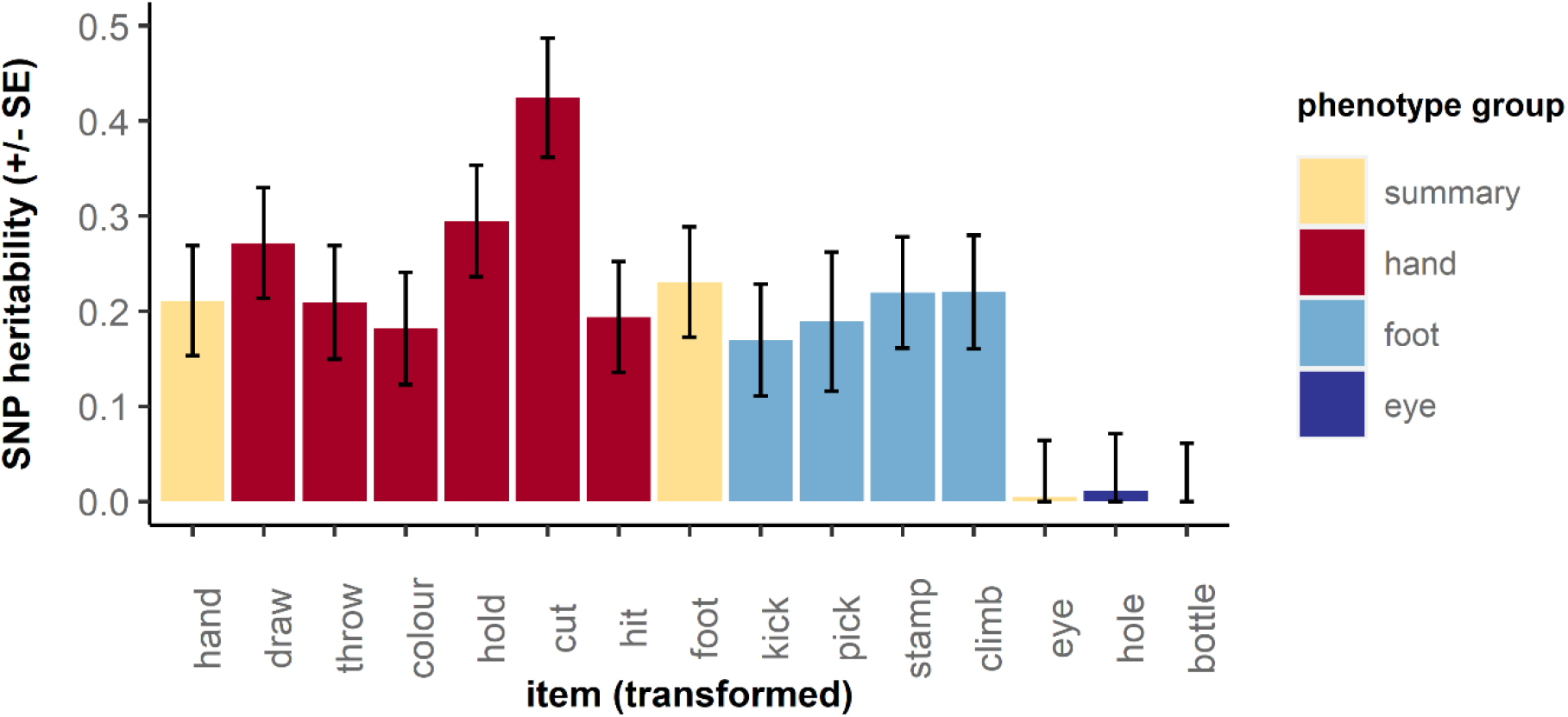
SNP-h^2^ estimates for laterality measures after transformation into quantitative scores in ALSPAC. Results are shown for individual items and summary measures (yellow). Bars represent standard errors.

For comparison, we estimated the SNP-h^2^ for the untransformed categorical items for left- and mixed-side preference categories. SNP-h^2^ for left-side preference ranged from .00 (*p* = .500) for “foot used to climb a step” to .13 (*p* = .031) for “hand used to cut” (Figure S8A, Table S14). SNP-h^2^ for mixed-side preference ranged from .00 (*p* = .500) for the hand preference summary item to .12 (*p* = .031) for “hand used to draw” (Figure S8B, Table S15).

Parent-offspring regression run on the transformed summary items suggested heritability estimates of .27 for handedness (95% CI = [.11, .42], *p* = 5.6 × 10^−4^), .09 for footedness (95% CI = [.01, .17], *p* = .030), and .08 for eyedness (95% CI = [-.04, .20], *p* = .198).

Univariate SEM analysis in the Hong Kong cohort resulted in heritability estimates of .45 for handedness (95% CI = [.29, .63]), .08 for footedness (95% CI = [.00, .25]), and .08 for eyedness (95% CI = [.00, .26]). Therefore, the heritability estimates for the quantitative phenotypes were consistently higher than for categorical measures, both for SNP-h^2^, parent-offspring, and twin SEM estimates.

### SEM

Multivariate GRM-SEM analysis was performed on the transformed R-M-L summary items for handedness, footedness, and eyedness in ALSPAC (Figure 3A). The squared path coefficient of genetic factor A1 explains genetic variance in hand preference (a_11_) and genetic variance that is shared with foot (a_21_) and eye preference (a_31_). A single genetic factor (A1) explained 20.36% of the phenotypic variance in handedness (a_11_ = 0.45, *p* = 2.4 × 10^−12^), 22.12% of the variance in footedness (a_21_ = 0.47, *p* = 9.2 × 10^−10^) and 3.84% of the variance in eyedness (a_31_ = 0.20, *p* = 9.2 × 10^−3^). All other path coefficients were non-significant, suggesting that one shared genetic factor (A1) contributes to a general left/right directionality across all three phenotypes.

**Figure 3:**
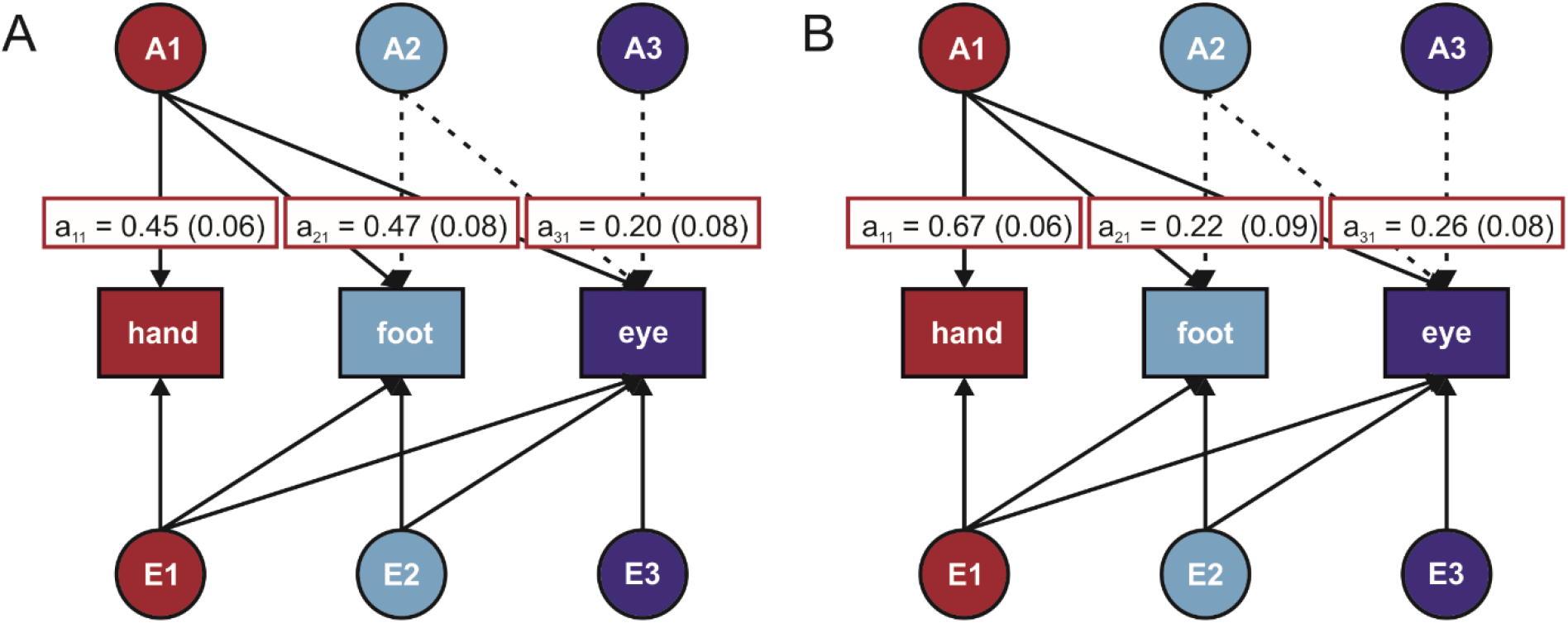
Results of SEM analyses between laterality phenotypes. A) Results of GRM-SEM in ALSPAC. B) Results of behavioural SEM in the Hong Kong cohort. Circles on top and bottom indicate genetic (A) and environmental (E) factors, respectively. Coloured boxes indicate the phenotypes. Solid lines indicate significant path coefficients, dotted lines indicate non-significant path coefficients. White boxes indicate path coefficients and standard errors (SE) for significant genetic factors. The contour of the white boxes indicates the genetic factor (A1 in all cases).

Bivariate heritability analysis confirmed that shared genetic influences accounted for 36.7% of the phenotypic correlation between handedness and footedness (*p* = 6.6 × 10^−6^), 24.9% of the correlation of between footedness and eyedness (*p* = .020), and 26.2% of the correlation between handedness and eyedness (*p* = .020). We replicated these findings with multivariate behavioural SEM in an independent cohort (*n* = 358). In the Hong Kong cohort, A1 explained 44.30% (95% CI = [28.50, 62.30]) of the phenotypic variance in handedness (a_11_ = 0.67, *p* < .001), 5.00% (95% CI = [0.20, 15.30]) of the variance in footedness (a_21_ = 0.22, *p* = .014), and 7.00% (95% CI = [0.80, 18.20]) of the variance in eyedness (a_31_ = 0.26, *p* = .003) (Figure 3B). All other path coefficients were non-significant, consistent with results for ALSPAC.

### PRS analysis

None of the PRS associations survived correction for multiple comparisons. The strongest association was found for PRS for IQ, suggesting that genetic predisposition towards higher IQ is associated with a tendency towards right-handedness (ß = -1159.21, SE = 414.71, PRS R^2^ = 0.13%, *p* = .005). PRS results for all *p*-value thresholds are reported in Table S16.

## Discussion

We investigated the heritability of hand, foot, and eye preference using multiple approaches. To the best of our knowledge, this is the largest study conducted to date for multiple laterality measures in the same individuals. Our analysis of family trios showed that the probability of being left-sided increased for any left-sided parent on the same trait, with stronger effects for hand and foot, rather than eye preference, in line with previous reports ^15,34^. Stronger maternal than paternal effects have been reported in studies focussing mainly on handedness ^20,23^. In ALSPAC, we found a stronger maternal than paternal effect for foot, but not hand or eye preference. This stronger maternal effect was detected in the whole sample (*n* = 4 960 trios) and confirmed in the subset with genetically confirmed paternity (*n* = 1 150 trios). Maternal/paternal effects could be explained with sex-linked genetic or parent-of-origin effects. For example, the imprinted *LRRTM1* gene was found to be associated with handedness under a parent of origin effect ^58^. Parent of origin effects might be more wide-spread than appreciated, but their detection requires family samples as opposed to the most commonly used singleton cohorts ^59^. Few examples of parent-of-origin effects have been reported, for example for language-related measures ^60–62^. Besides non-paternity, the reliability of the maternal report on laterality phenotypes could have affected our analysis. We confirmed strong correlation (*r* = .95) between the preferred hand for drawing assessed using maternal report at age 3.5 and self-reported preferred hand for writing in later childhood. The fact that more children switch hand preference from left to right ^63^ could indirectly suggest that switching attempts by parents or teachers have occurred at least until the mid 1990s. Overall, our analysis supports a genetic component underlying these laterality traits and highlights a specific maternal effect for footedness. The maternal effects could result from a higher genetic load required to manifest left-side preference in females. A similar buffering effect has been proposed to explain the higher prevalence of neurodevelopmental disorders in males ^64^.

Using transformed quantitative phenotypes ^48^, we estimated SNP-h^2^ for handedness, footedness, and eyedness to be .21, .23, and .00, respectively. The heritability estimate for handedness is similar to what has been reported in behavioural twin studies (h^2^ = .25) ^18,19^ but higher than observed in GWAS (SNP-h^2^ = .06) ^24,65,66^ for categorical handedness. Instead, estimates for categorical phenotypes were non-significant, suggesting that the transformed phenotypes are better suited to detect the genetic component underlying lateralised traits than binary phenotypes. Accordingly, behavioural analysis in the Hong Kong twin cohort revealed a heritability estimate of .45 for the quantitative handedness phenotype - much higher than what has been observed for a categorical measure of writing hand (.27) in the same cohort ^38^. Parent-offspring regression in ALSPAC also showed significant heritability for handedness and footedness when using the quantitative phenotypes. We conclude that the quantitative phenotypes are better suited to capture the polygenic nature of handedness as expected under a liability threshold model ^67^. The lack of association between the PRS derived from a recent large-scale GWAS for categorical handedness ^24^ suggests the influence of separate genetic factors. Lack of heritability for eyedness could reflect the poor quality of phenotype assessment, i.e. eyedness might be more difficult to assess and report accurately. Another possibility is that human eye preference does not have particular functional advantages and therefore the preferred side is less influenced by evolutionary forces and genetic factors. This is in contrast to other vertebrates such as bird ^68^ or fish species ^69^, where eye preference is involved in predator detection or social interaction.

Heritability estimates differed substantially between items used to assess handedness, footedness, and eyedness. We found the highest SNP-h^2^ for “hand used to cut” (with a knife). Previously, this item showed the weakest phenotypic correlation with the other questionnaire items ^70,71^ and the highest heritability ^33^. It has been proposed that summary items have reduced value to determine genetic factors involved in laterality ^32^. This was true for the handedness measure, but conversely, we observed higher SNP-h^2^ for the summary rather than single footedness items in ALSPAC, suggesting that in contrast to handedness, multiple items might better capture a genetic component for footedness. One possible interpretation is that multiple items will allow identifying mixed-footed rather than ambipedal individuals, who prefer both feet equally. Similar to Suzuki and Ando ^33^, our results suggest that the item “foot used to kick a ball”, which is often used as the only assessment item, is not the optimal choice to investigate the heritability of footedness. We previously showed that assessing footedness in terms of kicking systematically under-estimates the prevalence of mixed-footedness when compared to assessment using footedness inventories ^28^. Overall, there is no one correct measure for laterality items, however, our results demonstrate the importance of reporting data for single items ^72^ in addition to the aggregates and suggest the value of using multiple items.

All transformed items showed positive correlations on the phenotypic level. Previous research has shown a tendency towards a higher probability of left-sided lateral preferences in left-handers ^28,73^, suggesting that a common dimension of asymmetry underlies hand, foot, and eye preference ^74^. Multivariate SEM analysis supported the presence of one shared genetic factor explaining variance in handedness, footedness, and eyedness, but no unique genetic factors explaining independent variance for individual phenotypes in ALSPAC and the Hong Kong cohort. In ALSPAC, bivariate heritability analysis suggested that up to 37% of the phenotypic correlation is due to shared genetic effects.

An association between laterality and psychiatric disorders, especially schizophrenia ^75^, has long been debated. Of the different traits tested, we found suggestive evidence that PRS for IQ were associated with a tendency towards right-handedness, but not with footedness. Similarly, a recent dyslexia GWAS found positive genetic correlation between dyslexia and ambidexterity ^76^. A possible explanation for a specific link between cognitive measures and handedness is its association with language. It has been suggested that the higher prevalence of human right-than left-handedness has arisen from a left-hemispheric dominance for manual gestures that gradually incorporated vocalisation ^77^. Indeed, right-handers produce more right-than left-handed gestures when speaking ^78^. This would suggest that footedness and eyedness are phenotypically secondary to handedness, as has been suggested previously _79_.

## Conclusion

We assessed the heritability of multiple side preferences using family, genomic, and twin analyses. For footedness, stronger maternal than paternal effects highlight the necessity of examining parent-of-origin effects on the genetic level in future studies. SEM supports a shared genetic factor involved in all three phenotypes without independent genetic factors contributing to footedness and eyedness. The transformed quantitative phenotypes present a heritability that is higher than categorical measures in both molecular and behavioural analyses, suggesting that they might be better suited to identify the underlying genetic factors.

## Acknowledgments

We are extremely grateful to all the families who took part in this study, the midwives for their help in recruiting them, and the whole ALSPAC team, which includes interviewers, computer and laboratory technicians, clerical workers, research scientists, volunteers, managers, receptionists and nurses. We also thank Andrew P. Morris and Beate St Pourcain for advising on statistical analyses. The authors would also like to express their gratitude to Sarah Medland, Gabriel Cuellar Partida, and David Evans for providing the handedness GWAS summary statistics.

The UK Medical Research Council and Wellcome (Grant ref: 217065/Z/19/Z) and the University of Bristol provide core support for ALSPAC. This publication is the work of the authors and SP and JS will serve as guarantors for the analysis of the ALSPAC data presented in this paper. GWAS data were generated by Sample Logistics and Genotyping Facilities at Wellcome Sanger Institute and LabCorp (Laboratory Corporation of America) using support from 23andMe. Support to the genetic analysis was provided by the St Andrews Bioinformatics Unit funded by the Wellcome Trust [grant 105621/Z/14/Z]. The Hong Kong sample was funded through a Collaborative Research Fund from the Hong Kong Special Administrative Region Research Grants Council (CUHK8/CRF/13G, and C4054-17WF). JS is funded by the Deutsche Forschungsgemeinschaft (DFG, German Research Foundation, 418445085) and supported by the Wellcome Trust [Institutional Strategic Support fund, Grant number 204821/Z/16/Z]. For the purpose of open access, the author has applied a CC BY public copyright licence to any Author Accepted Manuscript version arising from this submission. SP is funded by the Royal Society (UF150663).

## Conflict of interest

The authors declare no competing financial interests in relation to the work described.

## Notes

### Competing Interest Statement

The authors have declared no competing interest.

