## Supplementary methods and figures for "Quantitative multidimensional phenotypes improve genetic analysis of laterality traits"

*Participants and phenotypes*

*ALSPAC:* Laterality phenotypes for children were assessed at 42 months based on maternal report. For mothers and fathers, laterality was assessed as self-report. Hand preference was assessed using eleven items for mothers and fathers (writing, drawing, throwing a ball, holding a racket, cleaning teeth, cutting with a knife, hammering, striking a match, using an eraser, dealing cards, threading a needle) and six items for children (drawing, throwing, colouring, holding a toothbrush, cutting, hitting). Foot preference was assessed using four items for mothers and fathers (kicking a ball, picking up a pebble, stepping on an insect, stepping onto a chair) as well as children with slightly different wording (kicking a ball, picking up a stone, stamping on something, climbing a step). Eye preference was assessed using two items for mothers and fathers (looking through a telescope, looking into a dark bottle) as well as children (looking through a hole, looking through a bottle). All items were rated on a 3-point scale (coded as left = 1, either = 2, right = 3). A mean value was calculated from the number of non-missing responses for each participant. R-M-L (right-mixed-left) items for hand, foot, and eye preference were derived by recoding the mean variables with the following criteria: left when scoring < 1.50, mixed when scoring ≥ 1.5 and ≤ 2.6, right when scoring > 2.6. R-L (right-left) items for hand, foot, and eye preference were derived by recoding mean variables as left when scoring < 2 and right when scoring ≥ 2.

*Hong Kong:* Hand, foot, and eye preference were assessed using a modification of the Edinburgh Handedness Inventory (EHI) [1]. The questionnaire was translated into Chinese and included six hand preference items (writing, drawing, holding scissors, brushing teeth, chopsticks, spoon), one foot preference item ("Which foot do you prefer to kick with?") and one eye preference item ("Which eye do you prefer to see things?"). Self-reported responses were coded on a 5 point scale (i.e. always left, usually left, no preference, usually right, always right). To make items comparable to ALSPAC, each item was recoded as 1 (usually left or always left), 2 (no preference) or 3 (usually right or always right). A mean score was generated for hand preference items that was recoded to a summary item according to the ALSPAC data (< 1.50: left, ≥ 1.50 and ≤ 2.60: mixed, > 2.60: right).

### Supplementary figures

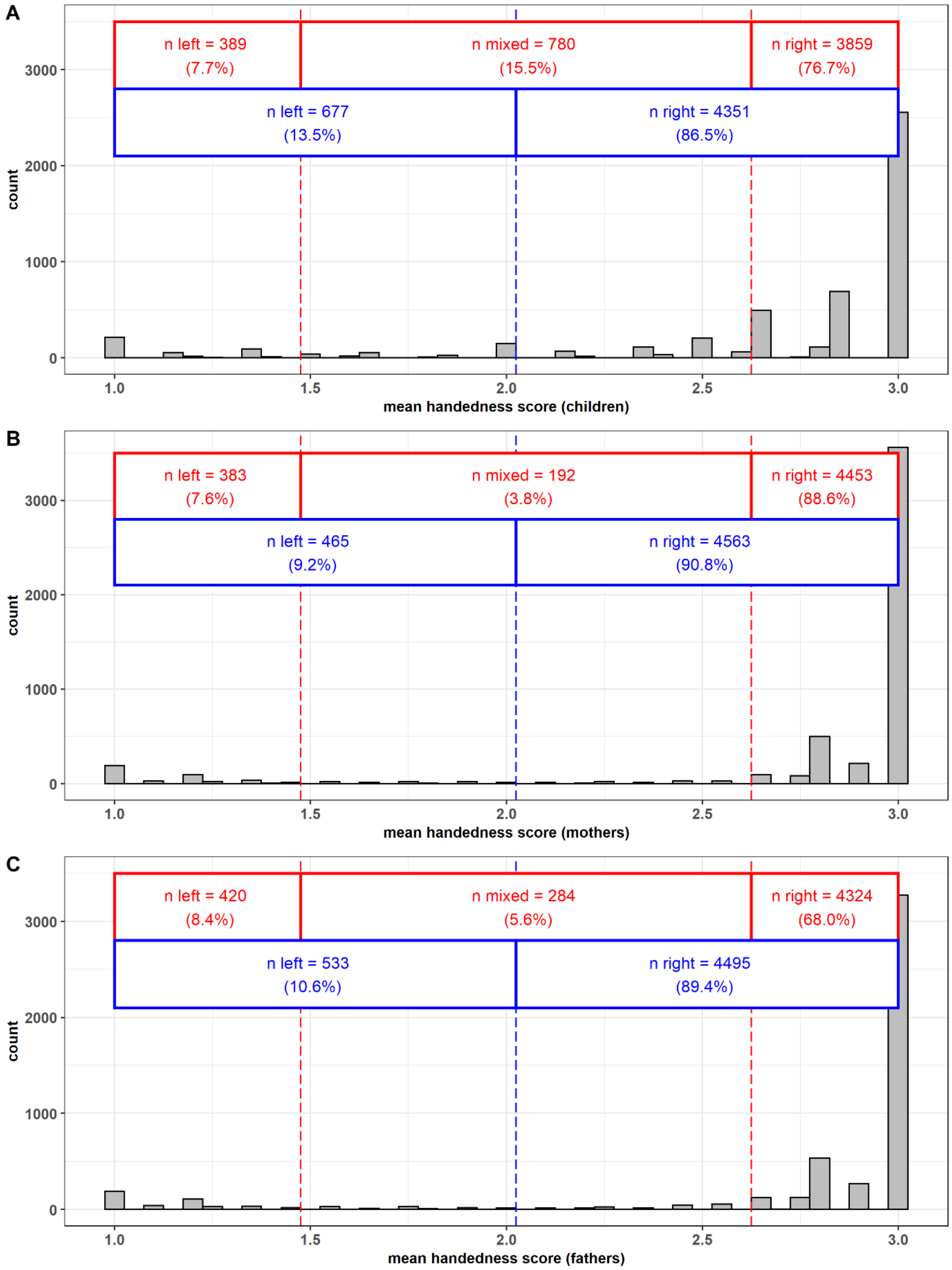


***Figure S1: Histograms showing the distribution of mean handedness scores for A) children, B) mothers, and C) fathers.*** *The red lines indicate the cutoff for the L-M-R summary items coded as left when < 1.5, mixed when > 1.5 and ≤ 2.6, and right when > 2.6. The blue lines indicate the cutoff for L-R with left ≤ 2 and right > 2. For each panel, the sample size is n = 5028.*

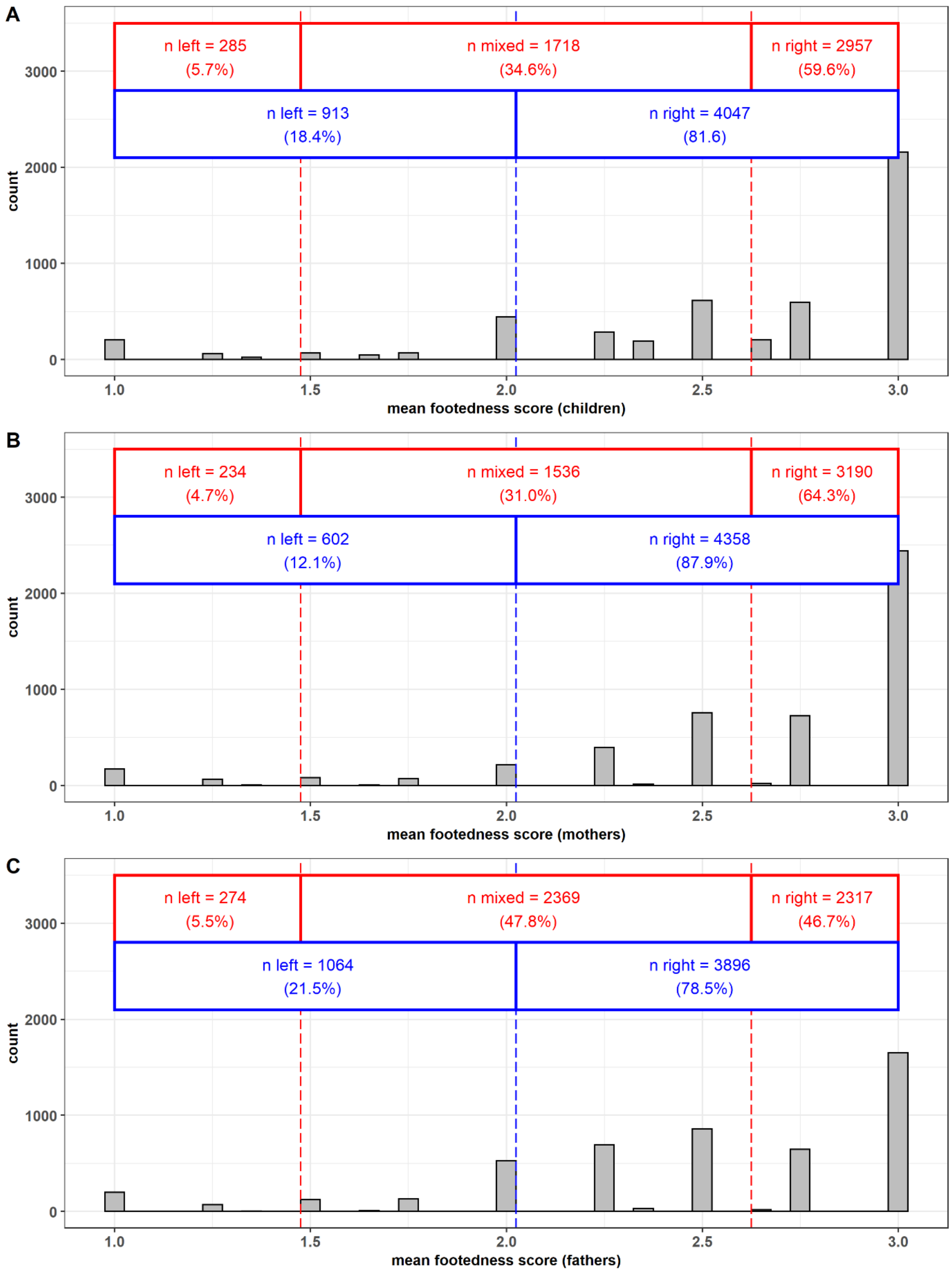


***Figure S2: Histograms showing the distribution of mean footedness scores for A) children, B) mothers, and C) fathers.*** *The red lines indicate the cutoff for the L-M-R summary items coded as left when < 1.5, mixed when > 1.5 and ≤ 2.6, and right when > 2.6. The blue lines indicate the cutoff for L-R with left ≤ 2 and right > 2. For each panel, the sample size is n = 4960.*

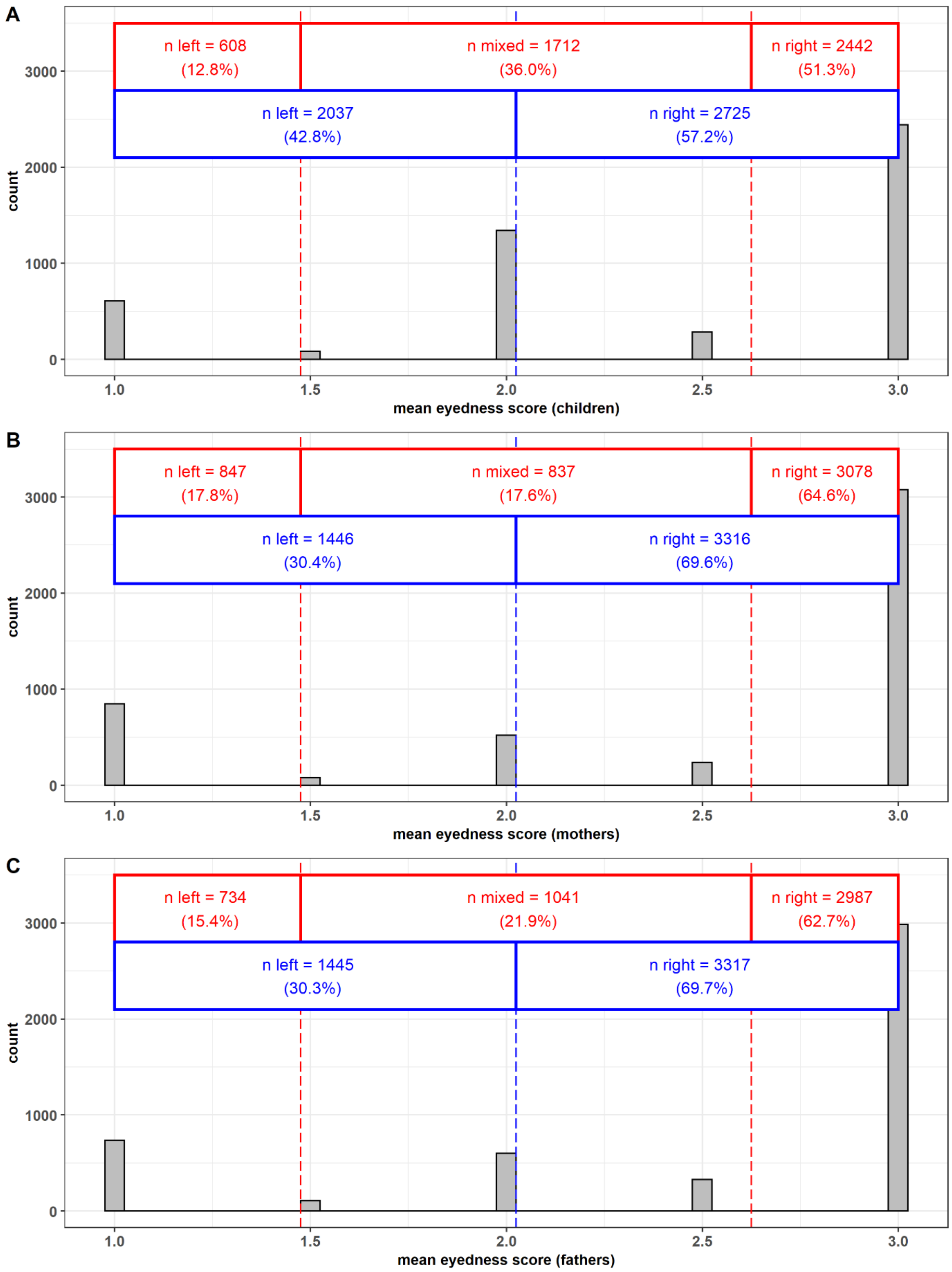


***Figure S3: Histograms showing the distribution of mean eyedness scores for A) children, B) mothers, and C) fathers.*** *The red lines indicate the cutoff for the L-M-R summary items coded as left when < 1.5, mixed when > 1.5 and ≤ 2.6, and right when > 2.6. The blue lines indicate the cutoff for L-R with left ≤ 2 and right > 2. For each panel, the sample size is n = 4762.*

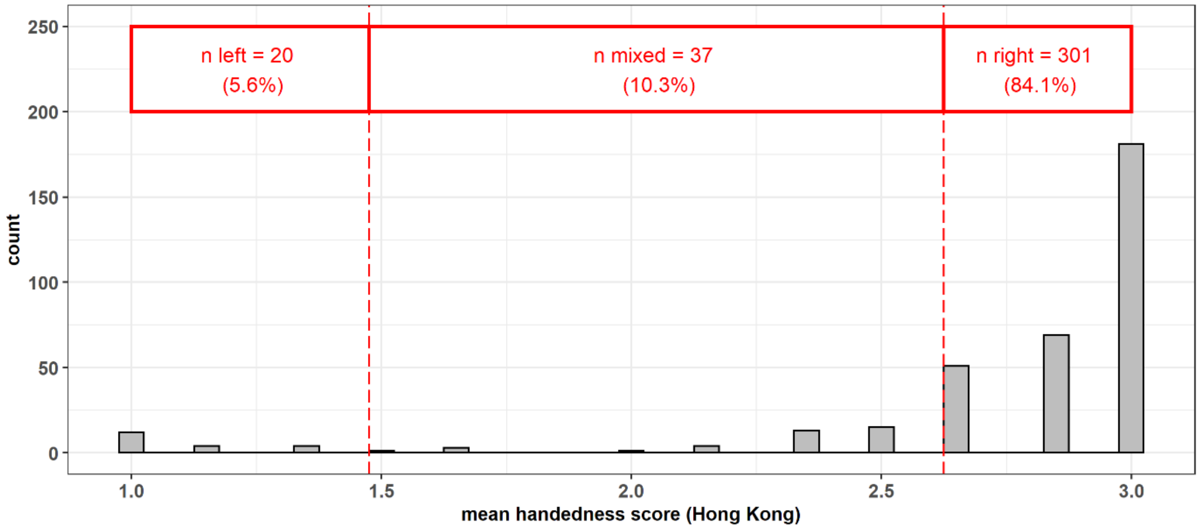


***Figure S4: Histogram showing the distribution of mean handedness scores in the Hong Kong cohort (n = 358).*** *The red lines indicate the cutoff for the L-M-R summary items coded as left when < 1.5, mixed when > 1.5 and ≤ 2.6, and right when > 2.6.*

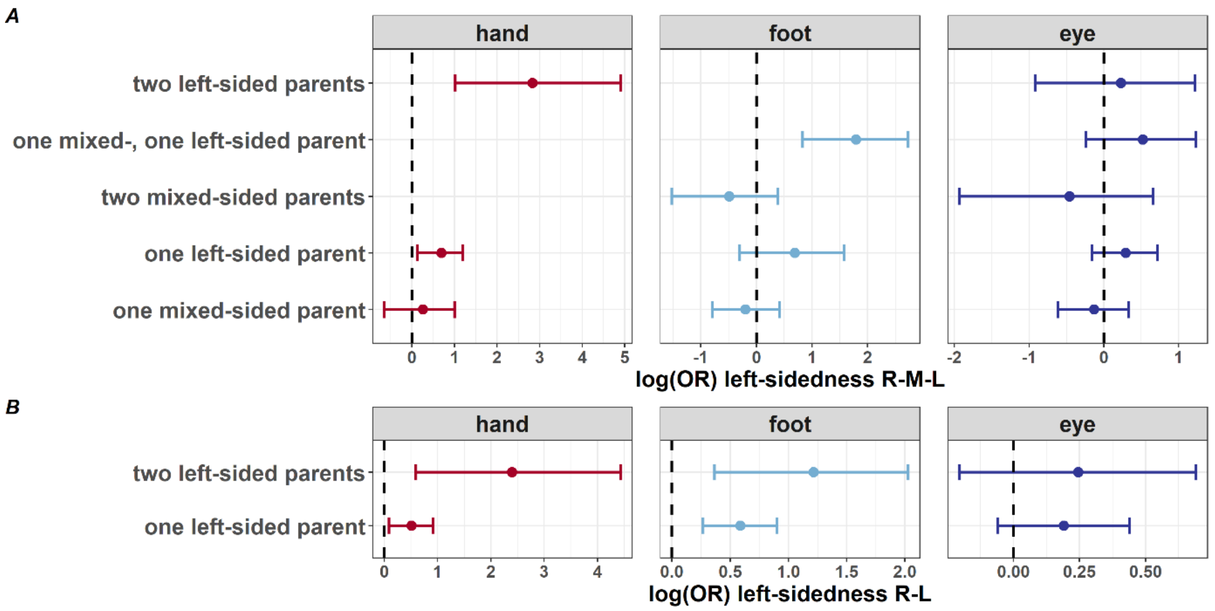


***Figure S5: Parental effects on child sidedness in the subset with confirmed paternity.*** *ORs [95% CI], resulting from logistic regression analysis.*

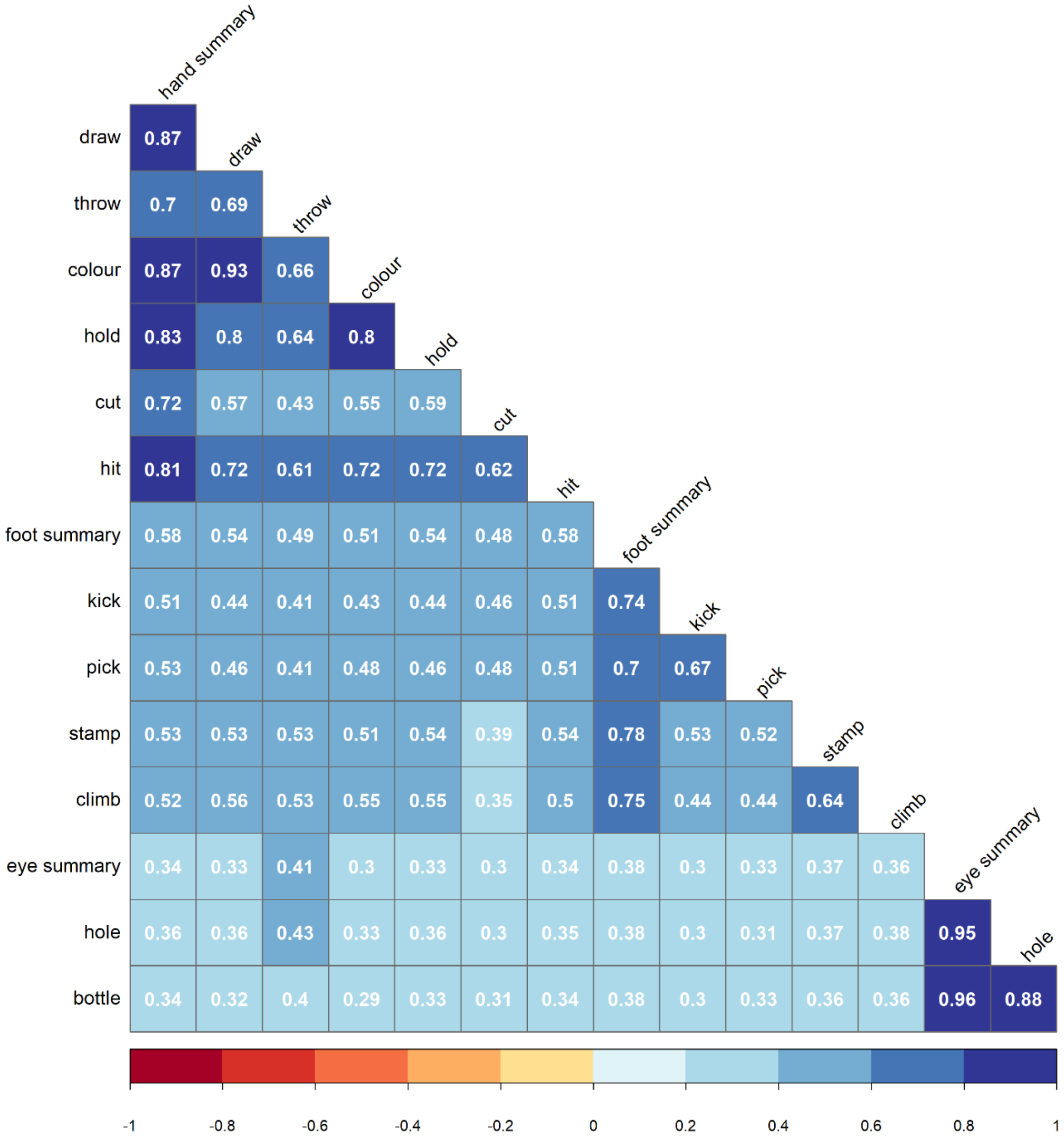


***Figure S6: Phenotypic correlations in the ALSPAC cohort.*** *Correlation coefficients are shown for the three summary items (handedness, footedness, and eyedness) and twelve single items after transformation (Pearson correlation). All correlations pass FDR correction for 105 comparisons.*

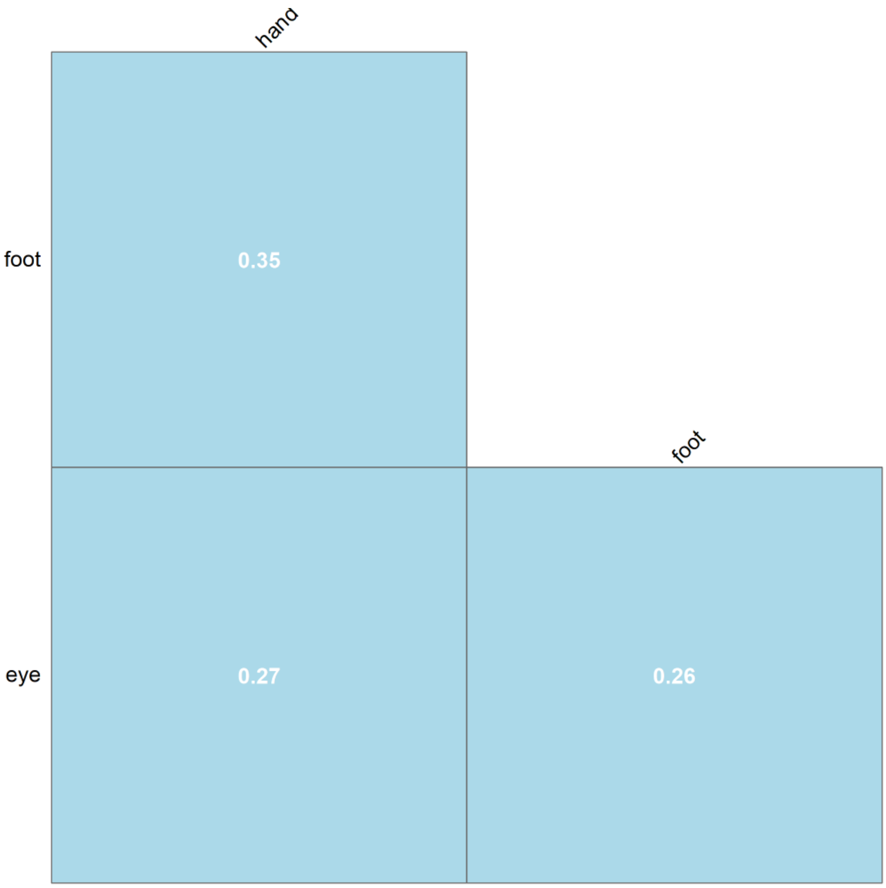


***Figure S7: Phenotypic correlations in the Hong Kong cohort.*** *Correlation coefficients are shown for the three items (handedness, footedness, and eyedness) after transformation (Pearson correlation). All correlations pass FDR correction.*

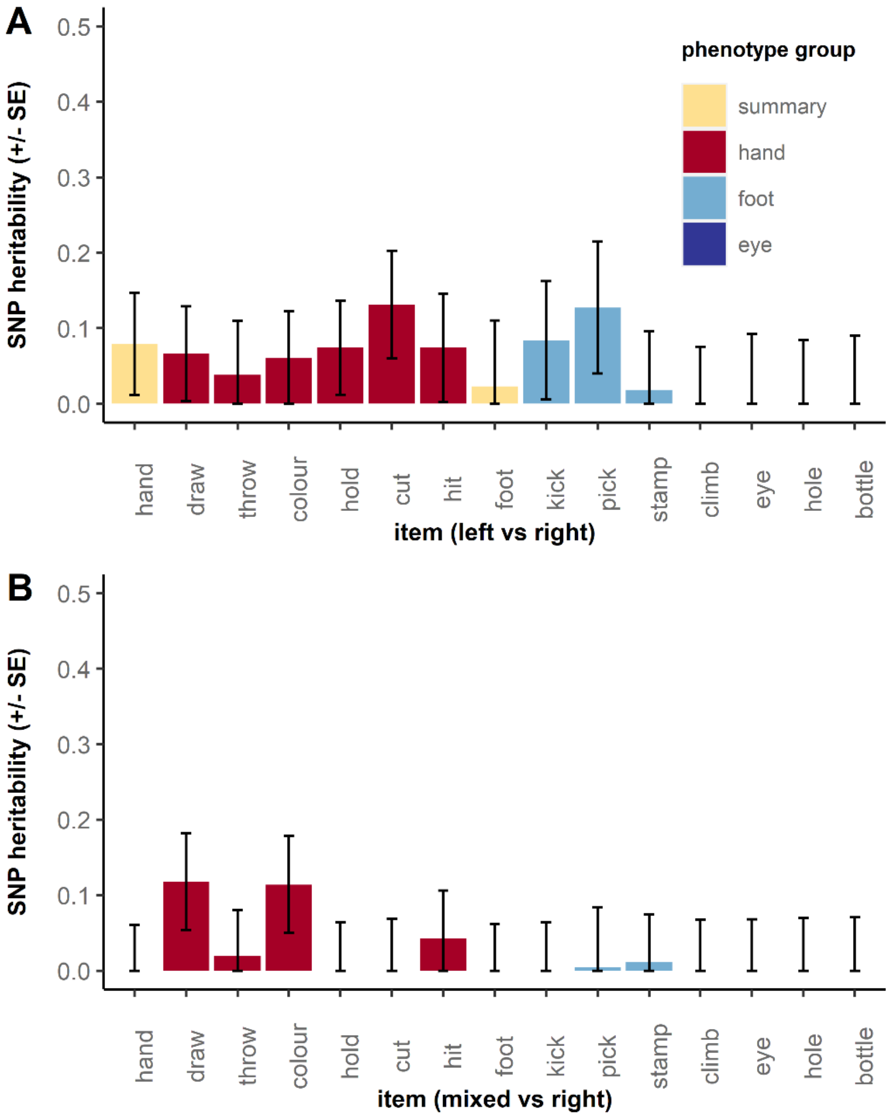


***Figure S8: SNP-$h^2$ estimates for untransformed laterality measures in ALSPAC.*** *Summary items for hand, foot and eye preference are shown in yellow, followed by the corresponding single items. The estimates were calculated using GCTA REML. Bars represent standard errors. A) Untransformed items (left vs. right), using sex, age, and two principal components as covariates. B) Untransformed items (mixed vs. right), using sex, age, and two principal components as covariates. Colour legend refers to what has been used in Figure 3 in the main text.*
